# First- and second-order contributions to depth perception in anti-correlated random dot stereograms

**DOI:** 10.1101/372169

**Authors:** Jordi M. Asher, Paul B. Hibbard

## Abstract

The binocular energy model of neural responses predicts that depth from binocular disparity might be perceived in the reversed direction when the contrast of dots presented to one eye is reversed. While reversed depth has been found using anti-correlated random-dot stereogram (ACRDS) the findings are inconsistent across studies. The mixed findings may be accounted for by the presence of a gap between the target and surround, or as a result of overlap of dots around the vertical edges of the stimuli. To test this, we assessed whether (1) the gap size (0, 19.2 or 38.4 arc min) (2) the correlation of dots or (3) the border orientation (circular target, or horizontal or vertical edge) affected the perception of depth. Reversed depth from ACRDS (circular no-gap condition) was seen by a minority of participants, but this effect reduced as the gap size increased. Depth was mostly perceived in the correct direction for ACRDS edge stimuli, with the effect increasing with the gap size. The inconsistency across conditions can be accounted for by the relative reliability of first- and second-order depth detection mechanisms, and the coarse spatial resolution of the latter.

## Introduction

Binocular depth perception depends on our ability to determine the difference in position of corresponding points between the two eyes’ images. This correspondence problem can be solved by using a matching measure to determine the image regions in each eye which are most similar in terms of the variation in local luminance intensity. This similarity matching can for example be based on the correlation of local intensity values.^1–4^ At the correct disparity offset between the left and right eyes, each point is expected to have similar luminance values in each eye, leading to a high interocular correlation. At incorrect disparities, non-corresponding points will be compared, which are likely to have different values of luminance, leading to low correlations.

In addition to the standard correlation-based measures described above, other matching-based metrics have been proposed that depend on detecting similarities between two image samples, based on the presence of individual matching features, rather than the overall correlation within the region.^5, 6^ These metrics are not necessarily mutually exclusive, and it has been suggested that both metrics are used independently and simultaneously.^5, 6^. Under the matching metric proposed, all evidence in favour of a match, when points have the same luminance polarity, contributes positively to the matching metric. However, in contrast to a standard correlation, points that have opposite contrast polarities, and provide evidence against a match, are ignored.

These similarity calculations can be related to the responses of binocular neurons in the visual cortex.^3, 4, 7, 8^ Neurons in V1 have a localised receptive field, consisting of both excitatory and inhibitory regions, and tend also to be tuned to orientation and spatial frequency.^3, 9–12^ Binocular neurons have a receptive field in both eyes, and their responses are thus sensitive to binocular disparities. Figure 1a shows an idealised binocular energy neuron,^3, 9, 10^ consisting of quadrature pairs of receptive fields for each eye. The responses of the first-stage filters are the square of the sum across the two eyes’ receptive field (figure 1b), which in turn are summed to computer the binocular energy response.

**Figure 1.**
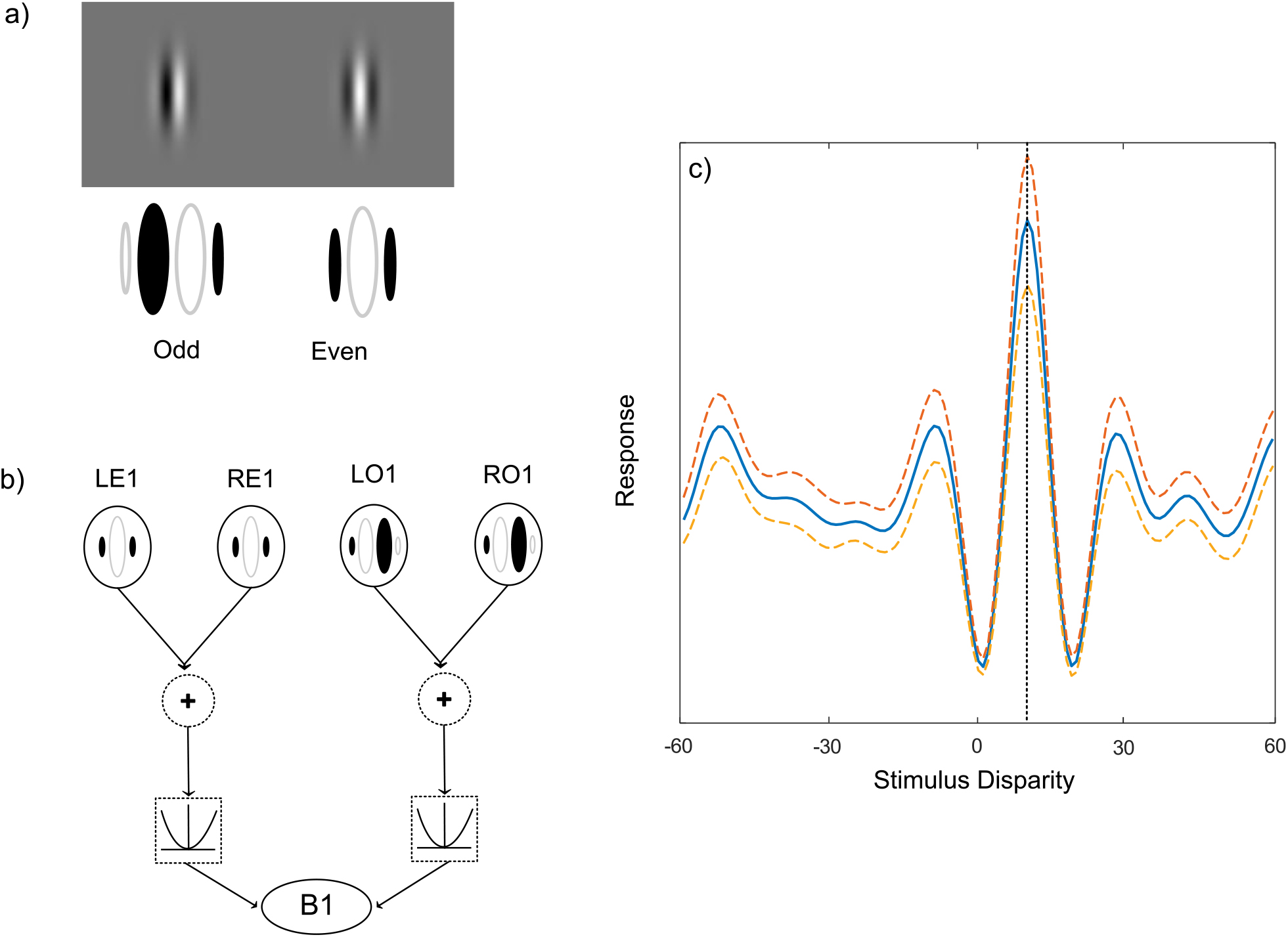
(a) Odd and even symmetric monocular receptive fields (b) These receptive fields are combined to compute the binocular energy response (c) The responses of binocular energy neurons show a characteristic tuning function. Here the response, calculated as the mean over 1000 Gaussian white noise samples, shows a peak at the disparity corresponding to the difference in position of the monocular receptive fields.

These components of the energy model have been used to characterise the responses of binocular simple and complex cells, respectively, although this simple hierarchy is best viewed as an idealisation of how these computations are performed.^13^ The response of a binocular energy neuron depends on the disparity in the images, forming a characteristic disparity tuning function (figure 1c). In this example, the receptive fields of the filters are shifted horizontally in the right eye compared to those in the left eye, meaning that this neuron responds most strongly to stimuli with the same disparity. This peak response at this optimum disparity is accompanied by disparities at which the response is reduced compared to baseline.

The binocular energy response is related to the point-wise correlation between the filtered left and right images.^3^ By summing across frequency, orientation and position,^3, 14, 15^ the correlation within a spatial neighbourhood can be calculated.^3, 4^ Complex cells in V1, as characterised by the binocular energy model, are thus well suited to support the calculation of the cross-correlation and other matching calculations thought to contribute to the solution of the correspondence problem.^9^

The dependence of the energy response on binocular cross-correlation means that manipulation of this correlation has provided a useful way of understanding how the responses of populations of binocular neurons contribute to the perception of depth.^2^ This is often investigated psychophysically using random-dot stereograms (RDS) which, by projecting an image that has been shifted slightly between the left and right eye, creates the perception of depth. Correlated random-dot stereograms (CRDS) present dots of the same luminance to each eye (Figure 2). However, an interesting application of RDS is the use of anti-correlated random-dot stereogram stimuli (ACRDS), where one eye’s view is replaced with its photographic negative.^5, 12, 16–23^ This means that the high positive correlations expected at the correct disparity become negative, and the disparity tuning function is inverted. Neurons in V1 show this inversion effect, but also a reduction in magnitude of their response that is not predicted by the energy model.^9–11^ This reduction in response has been modelled using the introduction of a threshold non-linearity,^24–26^ or a squaring of the energy response.^27^ These expansive non-linearities, by enhancing the difference between the amplitudes of the positive and negative peaks in the disparity tuning function, can be used to implement the cross-matching mechanism proposed by Doi and Fujita.^5, 6^

**Figure 2.**
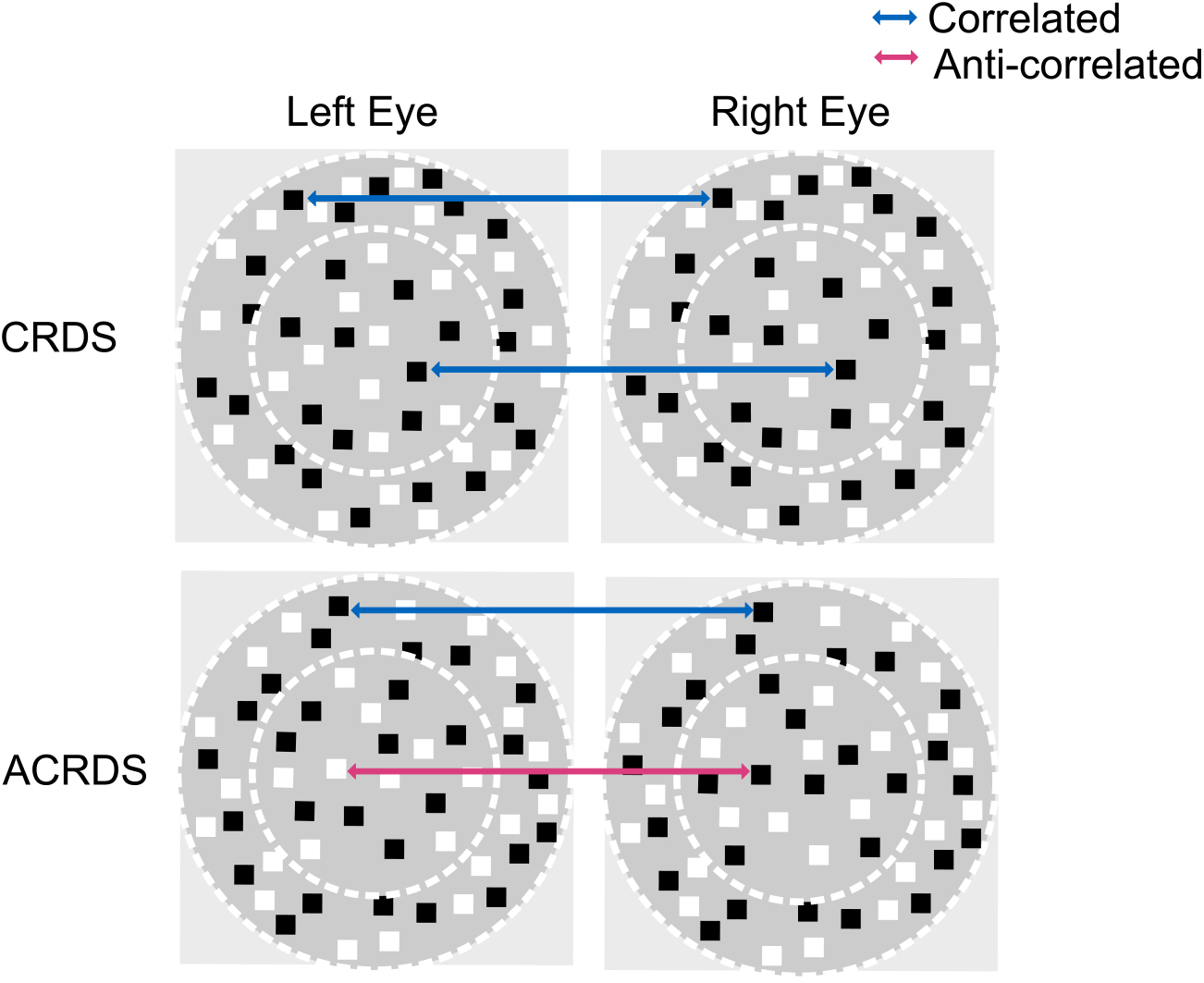
Correlated and Anti-correlated RDSs, left image is presented to the left eye and the right to the right eye. Both CRDS and ACRDS have a correlated reference (surround annulus). Correlated RDS (top) have correlated dots in the target (centre circle). In contrast that central target in anti-correlated RDS (bottom) have reversed luminance for the corresponding eyes.

In higher visual areas in the ventral stream, the responses of neurons tend not to be modulated by the disparity in ACRDS.^28, 29^ In contrast, neurons in dorsal stream areas show disparity tuning similar to that found in V1, but reduced in magnitude.^30^

In some psychophysical studies, the direction of depth perceived in ACRDS has been found to be reversed in comparison with equivalent CRDS.^5, 17, 21, 23^ One possible explanation of this percept is that it reflects the peaks in the inverted disparity tuning function, although whether these would signal a reversed or correct depth direction depends on the relationship between the spatial frequency and disparity tuning of the neuron, and the stimulus disparity (figure 3).

**Figure 3.**
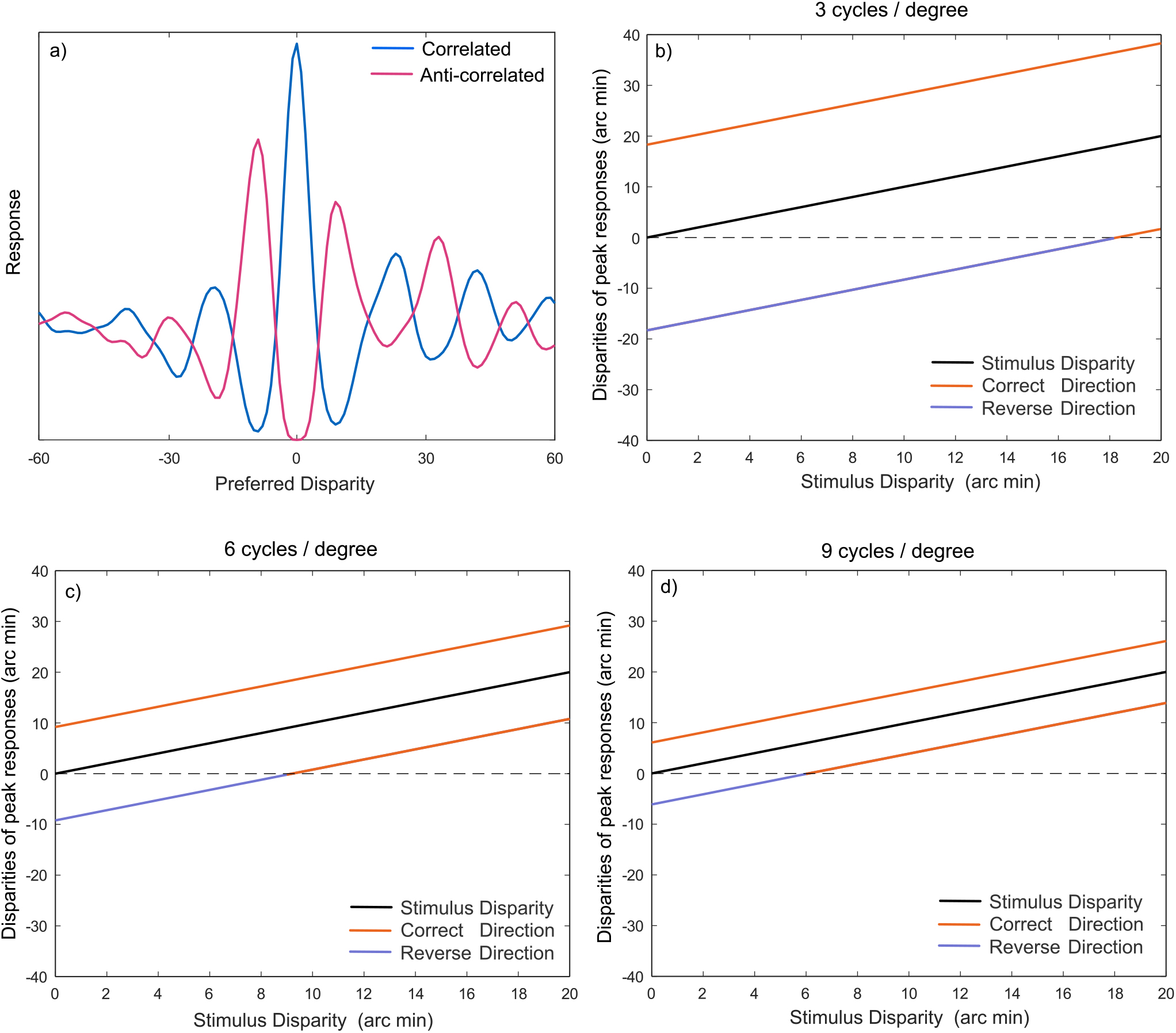
(a) An inverted disparity tuning function in response to an anti-correlated stereogram. (b-d) This can produce a peak at the opposite sign of disparity to that of the neuron’s preferred disparity. The sign of this false peak depends on the tuning frequency of the filter and the stimulus disparity. Each plot shows the peak of the disparity tuning function, as a function of disparity, averaged over 1000 Gaussian random noise stimuli. Invdividual plots show results for three different spatial frequencies

Alternatively, it has been suggested that the estimation of depth might reflect opponent processing, in which the difference between the responses of neurons tuned to equal but opposite disparities are calculated.^6, 27^ In this case, the negative correlations that exist in ACRDS would directly contribute to the reduction in the response to the correct disparity, thereby biasing towards the perception of depth towards the opposite direction.

Other studies have found no evidence for the perception of depth in ACRDS.^18, 19, 21^ This result is consistent with the fact that there is no coherent disparity, across different scales of analysis, in ACRDS, and also with the lack of disparity-selective responses in higher cortical areas.

The effect of decorrelation has been further assessed by Doi et al.^5^ who created stimuli containing an equal mixture of correlated and anti-correlated dot pairs. This results in an overall correlation of zero, so that a correlation-based mechanism predicts that depth would not be seen in these half-matched stimuli. In fact, depth is perceived in the correct direction. This is consistent with the responses of the cross-matching mechanism, which responds to the correct-matched dot pairs, but not to the anti-correlated pairs. Henriksen et al.^27^ showed that a similar prediction can be made by the squared energy response, which enhances the difference between the response to the paired and unpaired dots.

Predicting the perception of depth in ACRDS is complicated by the fact that there is no coherent disparity signal across different spatial scales. In CRDS, pooling of information across frequency, scale and position allows true peaks to emerge from amongst the many large responses that will occur at incorrect disparity values.^3, 14, 15^ This process does not produce a clear estimate for ACRDS, since large peaks are predicted to occur at different locations at different spatial scales.^19^

The perception of depth in ACRDS is further complicated by the existence of both first- and second-order mechanisms in stereoscopic processing. The discussion above considers the disparity information present in the Fourier components of the image, and how they might be combined. However, depth can also be perceived in second-order stimuli, containing informative disparities in variations in contrast, rather than in the underlying texture.^31^ Evidence for the existence of second-order channels has been provided by both psychophysical experiments, showing that participants can perceive depth in these stimuli,^31–41^ and physiological experiments, showing disparity-tuned responses to contrast envelopes.^14^ Tanaka and Ohzawa (2006)^14^ accounted for the responses of second-order neurons using a variation of the energy model. This model takes as its input not raw image values, but the outputs of monocular energy filters (figure 4). This monocular energy calculation is followed by a binocular energy calculation, with filters tuned to a much lower spatial frequency.

**Figure 4.**
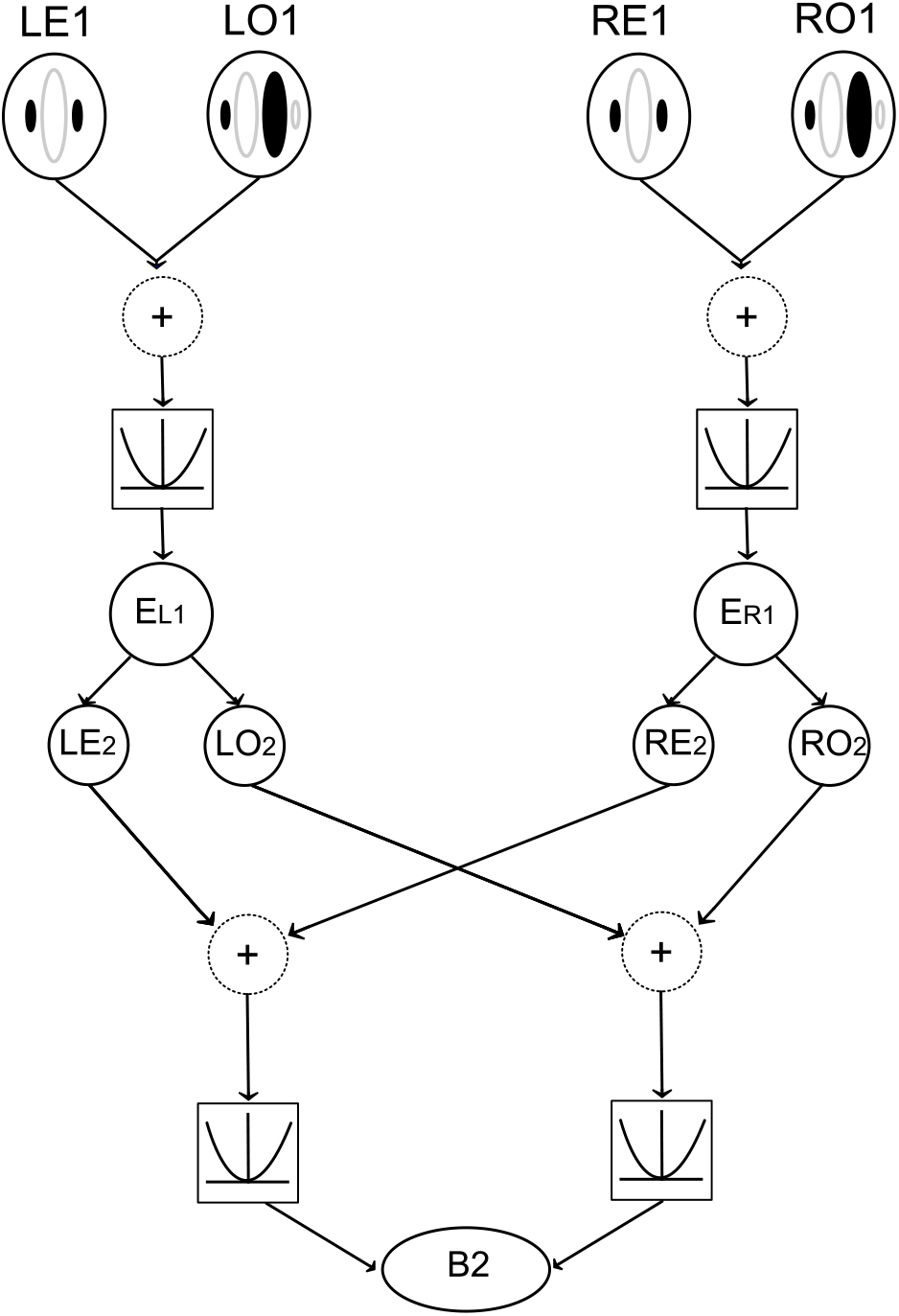
A second-order binocular energy model where monocular energy responses are calculated separately for each eye. These form the inputs into the second-order filters. Finally these are summed to calculate the second-order binocular energy response.

Wilcox and Hess^41^ showed that depth can be perceived from contrast envelopes even when the underlying random dot patterns are completely uncorrelated, provided that there was an informative disparity in the contrast variation. Sensitivity to depth from second-order stimuli is much poorer than that for first-order stimuli,^41^ and allows simple depth judgements to made, but not the perception of 3D shape.^42^ Second-order mechanisms will also provide disparity-tuned responses to reversed polarity stimuli, because the rectifying nonlinearity captures the magnitude, but not the contrast polarity, of luminance variations. An important distinction between first- and second-order channels is that the latter will signal depth in the forward, rather than reversed, direction. Cogan et al.^18^ proposed that depth from ACRDS relies on second-order mechanisms. In their experiments, they found forward depth perception for low density stimuli, but no reliable depth discrimination for high-density ACRDS.

The perception of depth in both CRDS and ACRDS also depends strongly on the presence of features at different disparities in the stimulus, such that the depth of a target can be judged based on its disparity relative to that at other locations. Large changes in absolute disparity, when they are not accompanied by changes in relative disparity, can go unnoticed by participants.^43–45^ Sensitivity to depth differences also falls as the spatial separation between target and reference increases.^46^ Karmihirata et al.^47^ argued that, due to the relatively weak disparity provided by ACRDS, depth is only perceived when a correlated reference is present, and there is no spatial gap between this and the anti-correlated target. Using stimuli in which the target was a circular region of anti-correlated dots, surrounded by an annulus of correlated dots, they showed that reversed depth was perceived when there was no-gap, but that this deteriorated when a small-gap was presented. The lack of a gap means that, once a non-zero disparity is incorporated into the stimuli, there will be overlap between the dots in the target and surround, such that both correlated and anti-correlated dots will fall into the receptive fields of neurons aligned with the vertical edges of the stimulus. This creates regions of decorrelation at the edges of the stimuli, on a different side in each eye. Decorrelation of this type occurs naturally through half-occlusion, whereby parts of a stimulus are visible to one eye but not the other (figure 5).^48–52^ Depth is perceived in these stimuli, consistent with this geometric interpretation. It is thus possible that the perception of depth in this case relates to the presence of decorrelation, although it should be noted that this would not explain depth discrimination in other cases^21^ in which there was a horizontal edge.

**Figure 5.**
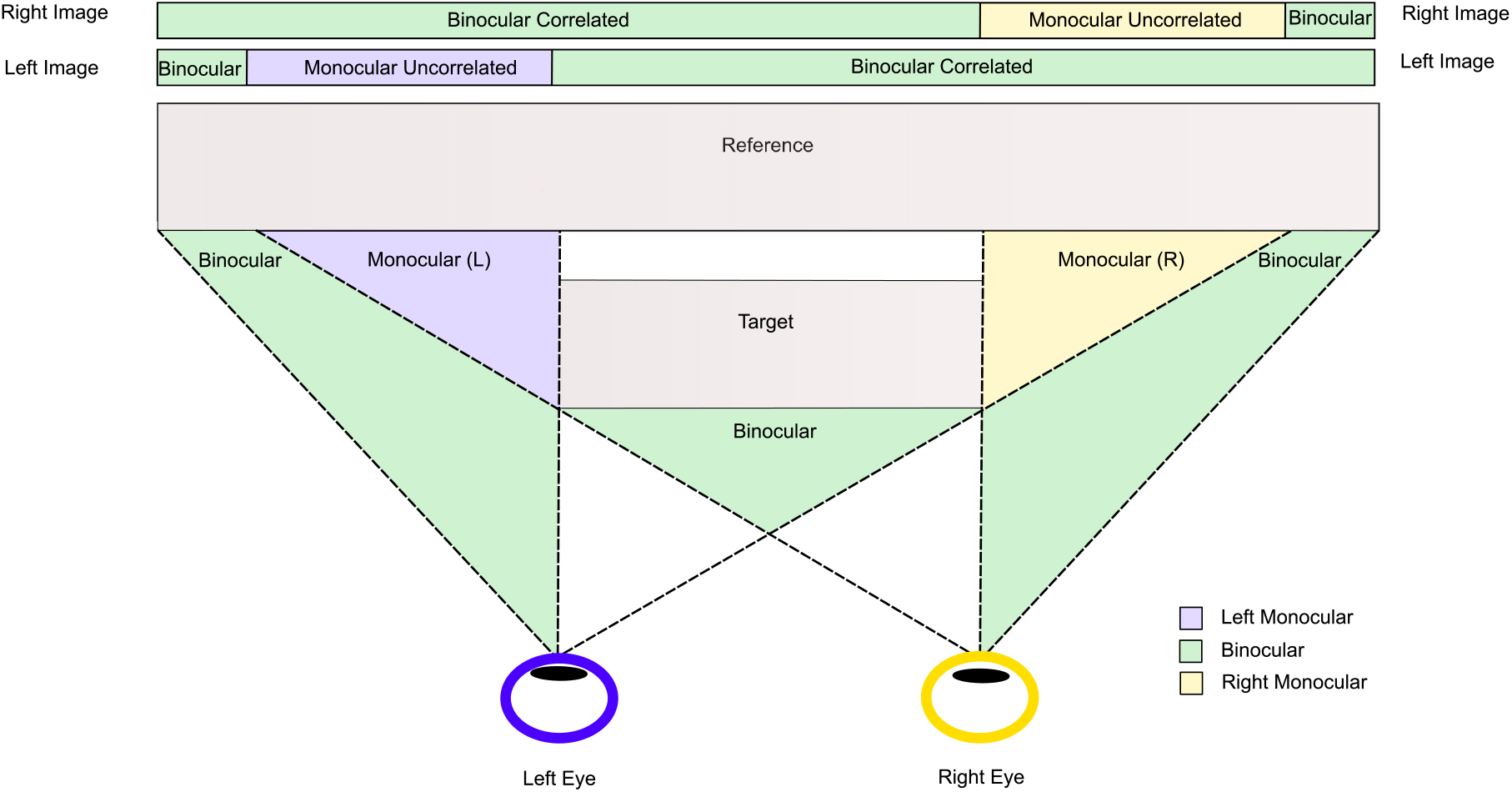
Schematic representation of the process whereby the different image for each eye can contain matched (correlated) and unmatched (uncorrelated) information

The first purpose of the current study was to understand the contribution of decorrelated edge regions, and the spatial separation between target and surround, on the perception of depth in ACRDS. This was done by using stimuli consisting of (i) an anti-correlated circular target surround by a correlated annulus (ii) a vertical edge between correlated and anti-correlated regions and (iii) a horizontal edge between correlated and anti-correlated regions. In the first two stimuli, there are regions containing both correlated and anti-correlated dots, whereas in the horizontal stimuli each region contains exclusively correlated or anti-correlated dots. The second purpose was to understand the contribution of mechanisms sensitive to first- and second-order disparities, and to monocular image regions, to the perception of depth in RDS. We did this firstly by modelling the responses of first- and second-order mechanisms to our stimuli, and secondly by manipulating the presence of decorrelated regions in our psychophysical experiments.

## Psychophysical Experiment

### Methods

#### Participants

10 participants (8 females, mean(std) age 24.5(9.6)) completed the experiment. All had normal or corrected to normal vision, and stereoacuity of at least 50 arc sec, as measured using Stereo Optical Butterfly Stereotest. All work was carried out in accordance with the Code of Ethics of the World Medical Association (Declaration of Helsinki). The study procedures were approved by the University of Essex University Ethics Committee (Application No. JA1601) All participants gave informed written consent and received payment for their participation.

#### Apparatus

Stimuli were presented on a VIEWPIXX 3D monitor, viewed from a distance of 40 cm. The monitor screen was 52 cm wide and 29 cm tall. The screen resolution was 1920 by 1080 pixels, with a refresh rate of 120 Hz. Each pixel subtended 2.4 arc min. Stereoscopic presentation was achieved using a 3DPixx IR emitter and NVIDIA 3D Vision LCD shutter glasses. The cross-talk between the left and right images, measured using a Minolta LS-110 photometer, was 0.12%. Participants’ responses were recorded using the computer keyboard. Stimuli were generated and presented using MATLAB and the Psychophysics Toolbox extensions.^53–55^

#### Stimuli

Stimuli in all conditions were random dot stereograms, comprised of 12 arc min square red (27.0*cdm^−^*^2^) and black (0*cdm^−^*^2^) dots against a red (13.5*cdm^−^*^2^) background. Equal numbers of red and black dots were presented, with a total density of 1.12 dots/degree^*−*2^. In all cases, stimuli consisted of a correlated reference region, presented with 0 disparity, and a test region which was either correlated or anti-correlated. Stimuli were presented in three conditions: circular, horizontal and vertical (Figure 6).

**Figure 6.**
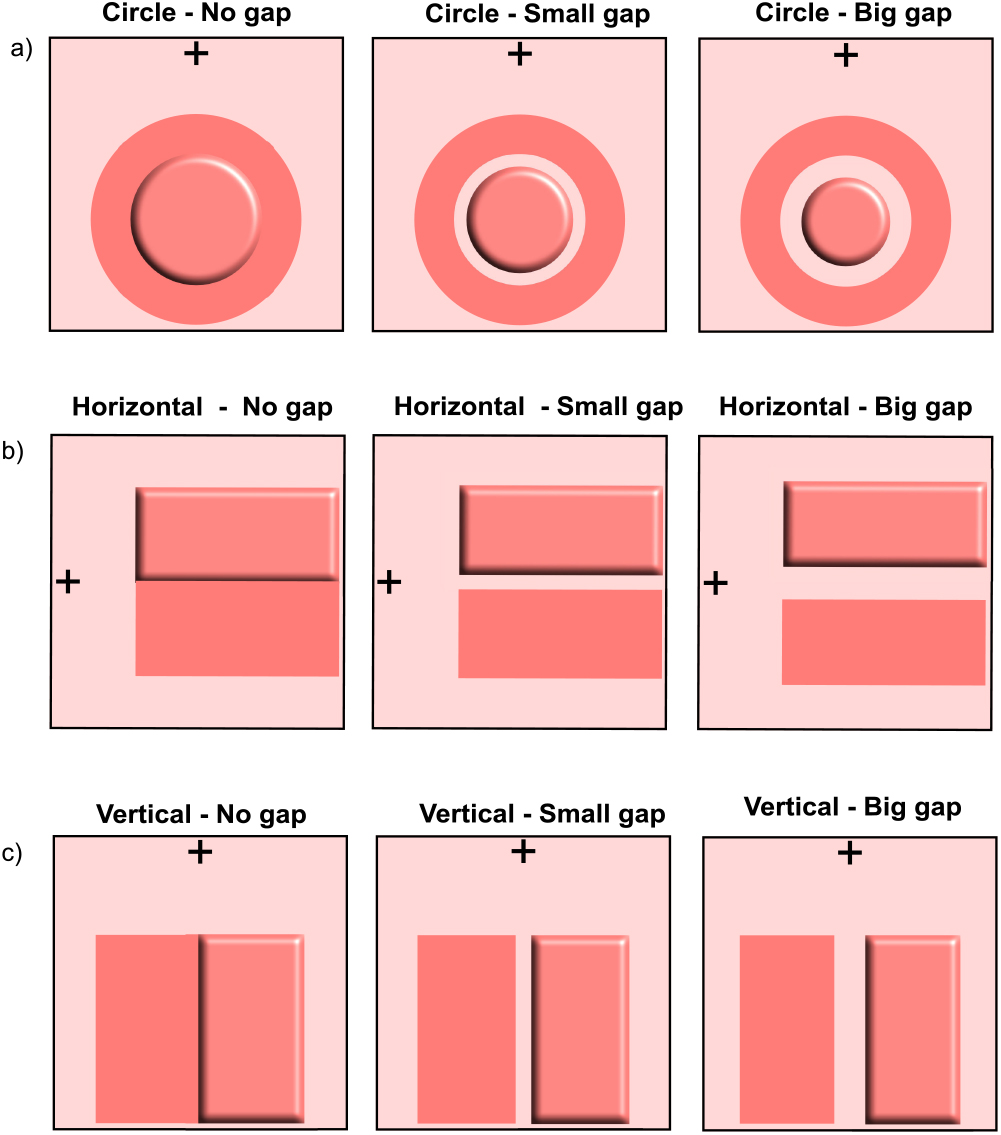
Schematic of stimuli used for each condition. a) For the Circle condition, the target was always the inside circle. b) For the Horizontal condition the target was always the right rectangle and c) for the Vertical condition the target was always the top rectangle. The task in each case was to identify whether the target is behind or in front of the reference. Participants were required to stay fixated on the cross at all times. In this schematic the target is always in front of the reference.

**Figure 7.**
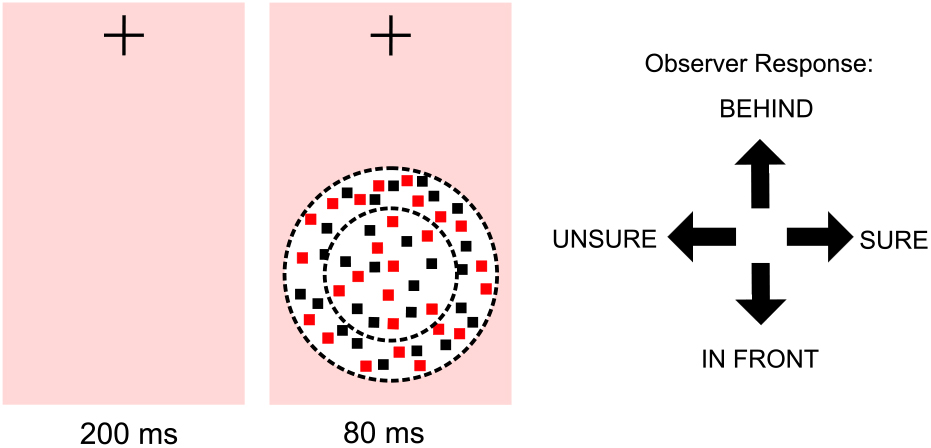
Participants were required to indicate for each presentation whether the target was in front of or behind the reference (surround). Immediately following the depth response a confidence decision (sure or unsure) was required.

#### Circular

The target was a circular region with a diameter of 4.4 degrees. The reference was a surrounding annulus, with no-gap, a small-gap (19.2 arc min) or a large gap (38.4 arc min) between target and the inside of the reference region (figure 6a). The inner diameter of the reference annulus depended on the size of the gap between the test and reference, and the outer diameter was 1.83 degrees larger than the inner radius. The circular stimuli were presented with the centre of the test region 5.5 degrees below the fixation cross.

#### Horizontal Edge

The reference and test regions were both a rectangle with a width of 5.5 degrees and a height of 2.75 degrees. Both were centered 5.5 degrees to the right of fixation, with the test presented above and the reference presented below the fixation cross. There were three levels of vertical gap; no-gap, a small-gap (19.2 arc min) or a large gap (38.4 arc min) between the reference and test regions (figure 6b).

#### Vertical Edge

The reference and test regions were both a rectangle with a width of 2.75 degrees and a height of 5.5 degrees. Both were centred 5.5 degrees below the fixation cross, with the test presented to the left and the reference presented to the right. As with the previous conditions, there was a horizontal separation of either no-gap, a small-gap (19.2 arc min) or a large gap (38.4 arc min) between the reference and test regions (figure 6c).

The dots were in all cases randomly repositioned on each frame. The test region was presented with a disparities of *±* 5.5, *±* 11 and *±* 22 arc min.

#### Procedure

Each trial began with the presentation of a central fixation cross for 200ms, followed by the presentation of a stimulus for 80ms. The fixation cross remained visible throughout each trial block.

After stimulus presentation, the participant was required to make two responses. The first was to indicate whether the target appeared closer (down arrow) or further way (up arrow) than the reference. The second was to indicate whether they felt confident in their response (right arrow) or that they were guessing (left arrow). The next trial began after the two responses had been made.

Each participant completed 18 blocks of trials, for all combinations of the three configurations, three separations, and two correlation conditions. In each block, each of the 6 disparities was presented 20 times. Block were presented in a randomised order, and separated over two or more sessions.

### Results

#### Depth Perception

To determine whether participants judgements shifted from ‘far’ to ‘near’ (or vice versa) as disparity shifted from uncrossed to crossed, results were analysed with a generalised linear mixed effects model. In this model, disparity was used as a predictor, with a probit linking function, and random slopes and intercepts across participants were included. A separate model was fit for each combination of shape, correlation and stimulus separation. This model allows us to determine whether, at the population level, forward or reversed depth was perceived, while also taking account of variation across participants.

Previous studies have found significant individual differences in depth perception in ACRDS, for example with some participants perceiving reversed depth but others perceiving no depth.^21^ We therefore fit the same generalised linear model, with a probit linking function, to the data for each participant separately. In each case, we then determined whether that participant perceived forward or negative depth for each stimulus based on whether the the slope parameter of the model has a significant positive or negative value. The results are shown in figure 11, which plots the number of participants perceiving forward depth, reversed depth, or no significant depth, for each stimulus type.

#### Circular Stimuli

The proportion of near judgements is plotted as a function of disparity. Results are plotted separately for CRDS and ACRDS and for the three separation distances. Figure 8 shows the mean results across participants. For CRDS, there was a positive slope for all separations indicating forward depth. In contrast there was no significant slope in either direction for ACRDS indicating no reliable depth perception.

**Figure 8.**
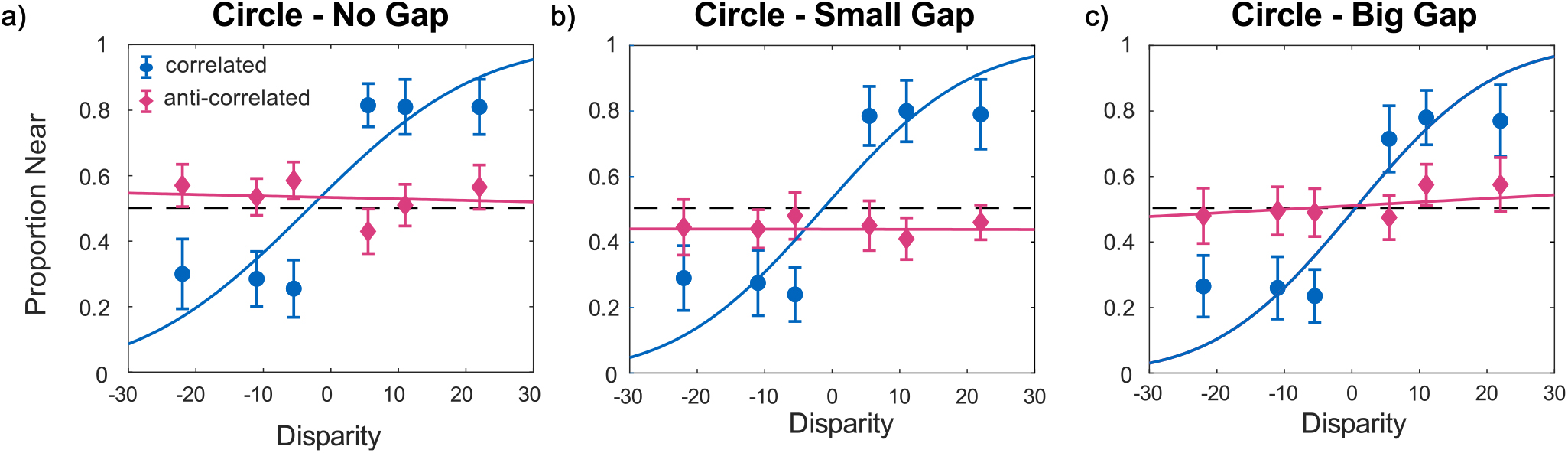
Mean proportion of near responses as a function of disparity for Circular stimuli for a) no-gap b) small-gap and c) big-gap. Separate plots chart the responses to anti-correlated stimuli (pink) and correlated responses (blue). The dashed line for each plot reflects performance at chance, uncrossed and crossed disparities are shown with negative and positive values. Error bars are *±*1 standard error of the mean.

Individual analyses showed that the majority of participants perceived forward depth for CRDS, while some were not able to reliably report the direction of depth for the larger stimulus separations. For CRDS, in most cases there was no reliable perception of depth. With no separation, 4/10 participants reported reversed depth, but no reversed depth was perceived for larger separations, consistent with recent findings.^47^ However, for the largest separation, there was evidence of reliable perceived depth in the forward direction for two participants.

#### Horizontal Stimuli

Data are presented in figure 9(a-c) in the same format for circular stimuli. For CRDS, there as a positive slope for all separations. For ACRDS, there was a significant positive slope for the small separation, consistent with forward, rather than reversed, depth.

**Figure 9.**
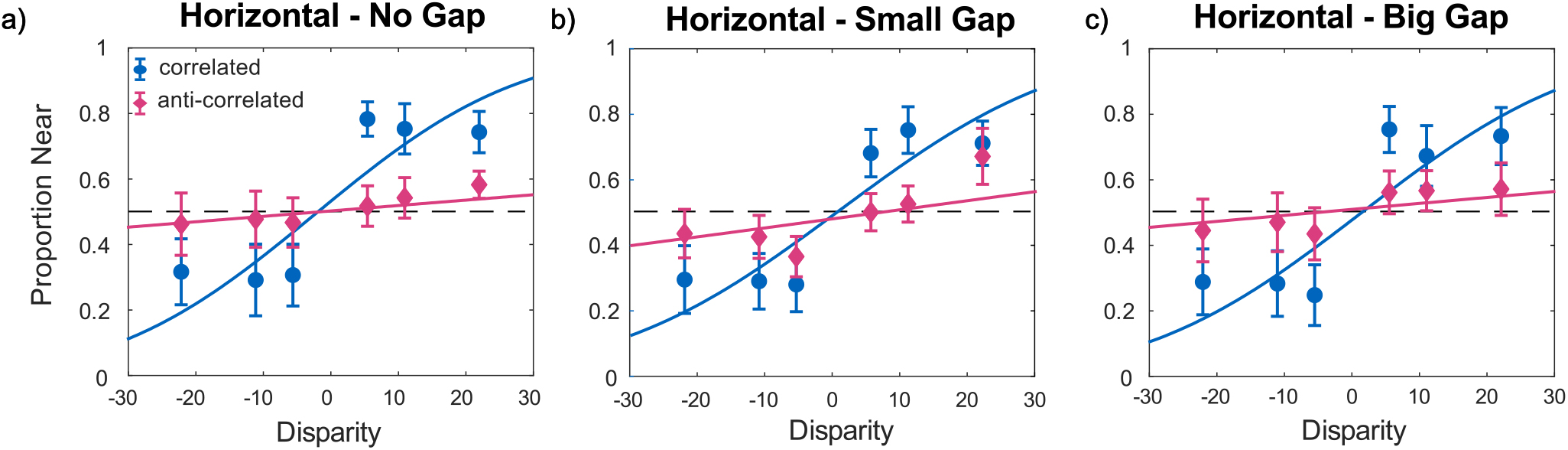
Mean proportion of near responses as a function of disparity for Horizontal stimuli for a) no-gap b) small-gap and c) big-gap. Separate plots chart the responses to anti-correlated stimuli (pink) and correlated responses (blue). The dashed line for each plot reflects performance at chance, uncrossed and crossed disparities are shown with negative and positive values. Error bars are ± standard error of the mean.

Individual analyses showed that the majority of participants perceived forward depth for CRDS, although this proportion decreased with increasing separation. For ACRDS, one participant perceived depth in the reverse direction when there was no-gap, and a minority of participants perceived depth in the forward direction.

#### Vertical Stimuli

Data for the vertical stimuli are presented in figure 10(a-c). For CRDS, there as a positive slope for all separations. For ACRDS, there was a significant positive slope for the two non-zero gaps, consistent with forward depth.

**Figure 10.**
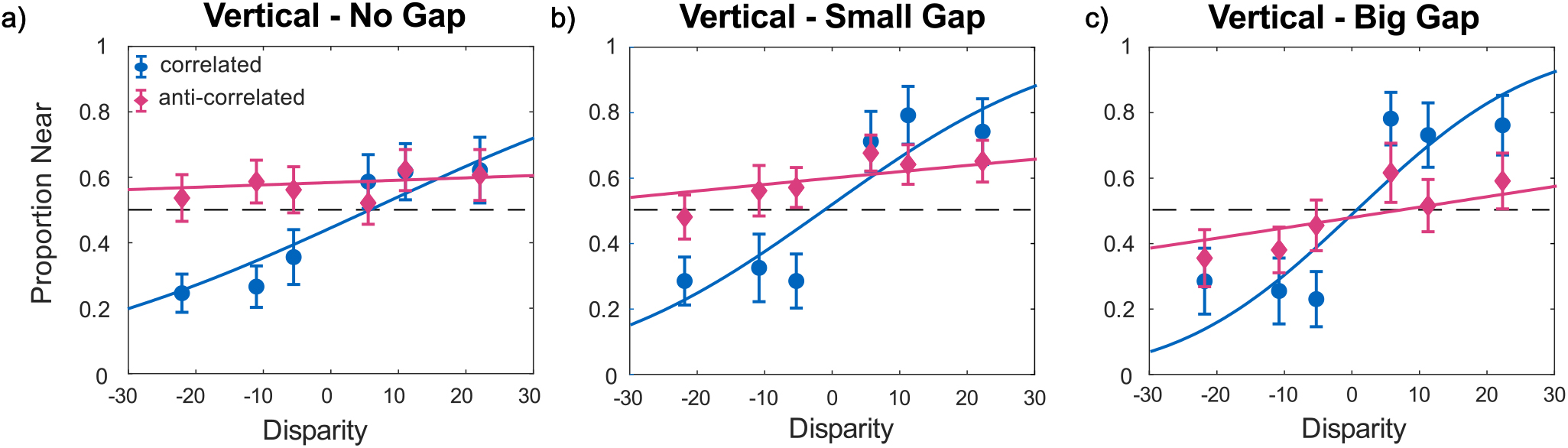
Mean proportion of near responses as a function of disparity for Vertical stimuli for a) no-gap b) small-gap and c) big-gap. Separate plots chart the responses to anti-correlated stimuli (pink) and correlated responses (blue). The dashed line for each plot reflects performance at chance, uncrossed and crossed disparities are shown with negative and positive values. Error bars are *±*1 standard error of the mean.

**Figure 11.**
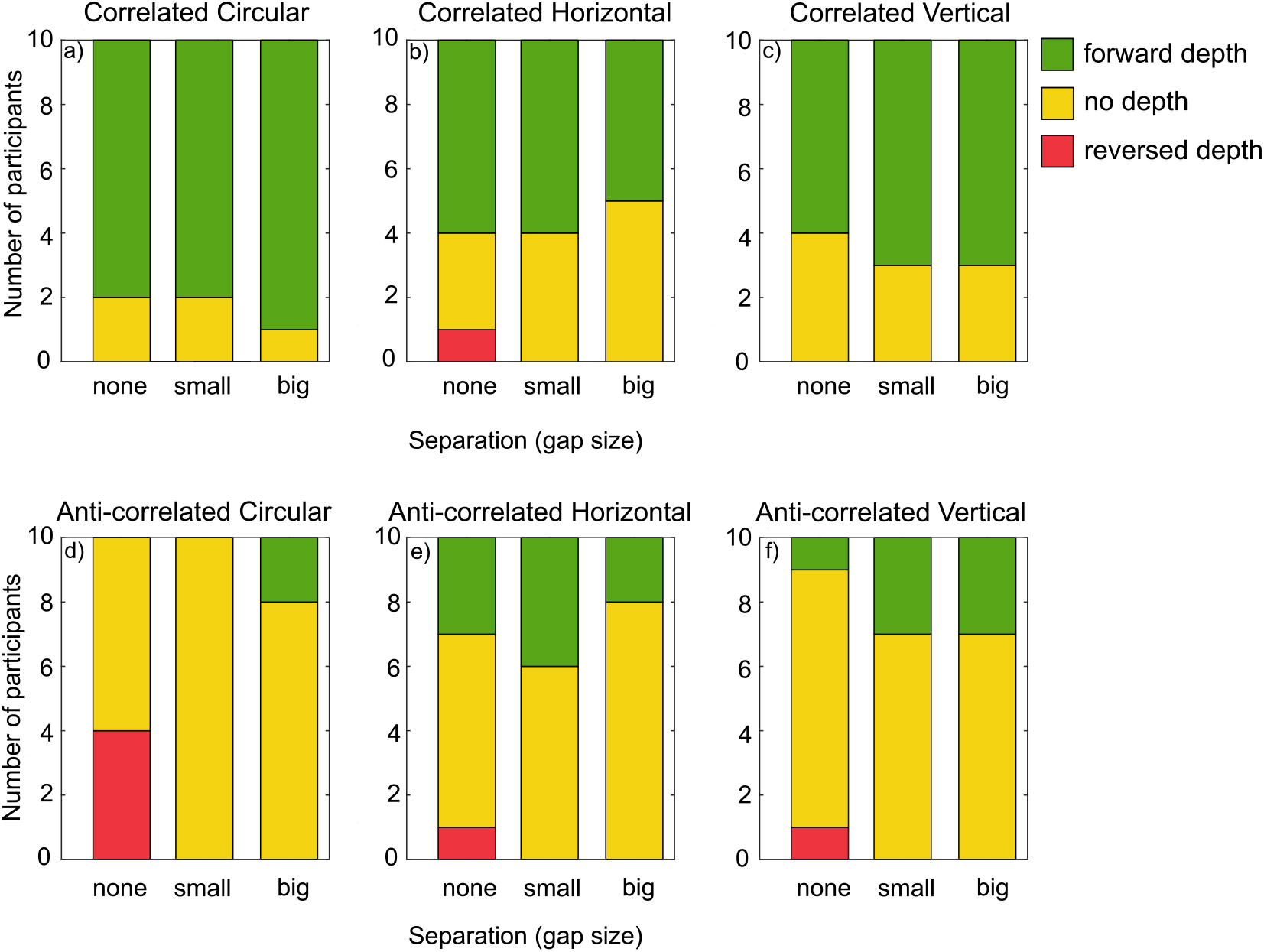
The proportion of participants reporting forward (green) reversed (red) or no (yellow) depth

Individual analyses showed very similar results to those found with a horizontal edge. The majority of participants perceived forward depth for CRDS, and this proportion decreased with increasing separation. For ACRDS, the majority of participants did not reliably perceive depth, but a minority did consistently perceive forward depth, and there was only one example of reliable reversed depth.

#### Confidence

Mean confidence ratings are plotted as a function of disparity for each combination of shape, correlation and distance in figure 12. Results are averaged across signs of disparity since, unlike depth judgements, we do not expect opposite results for near and far stimuli.

**Figure 12.**
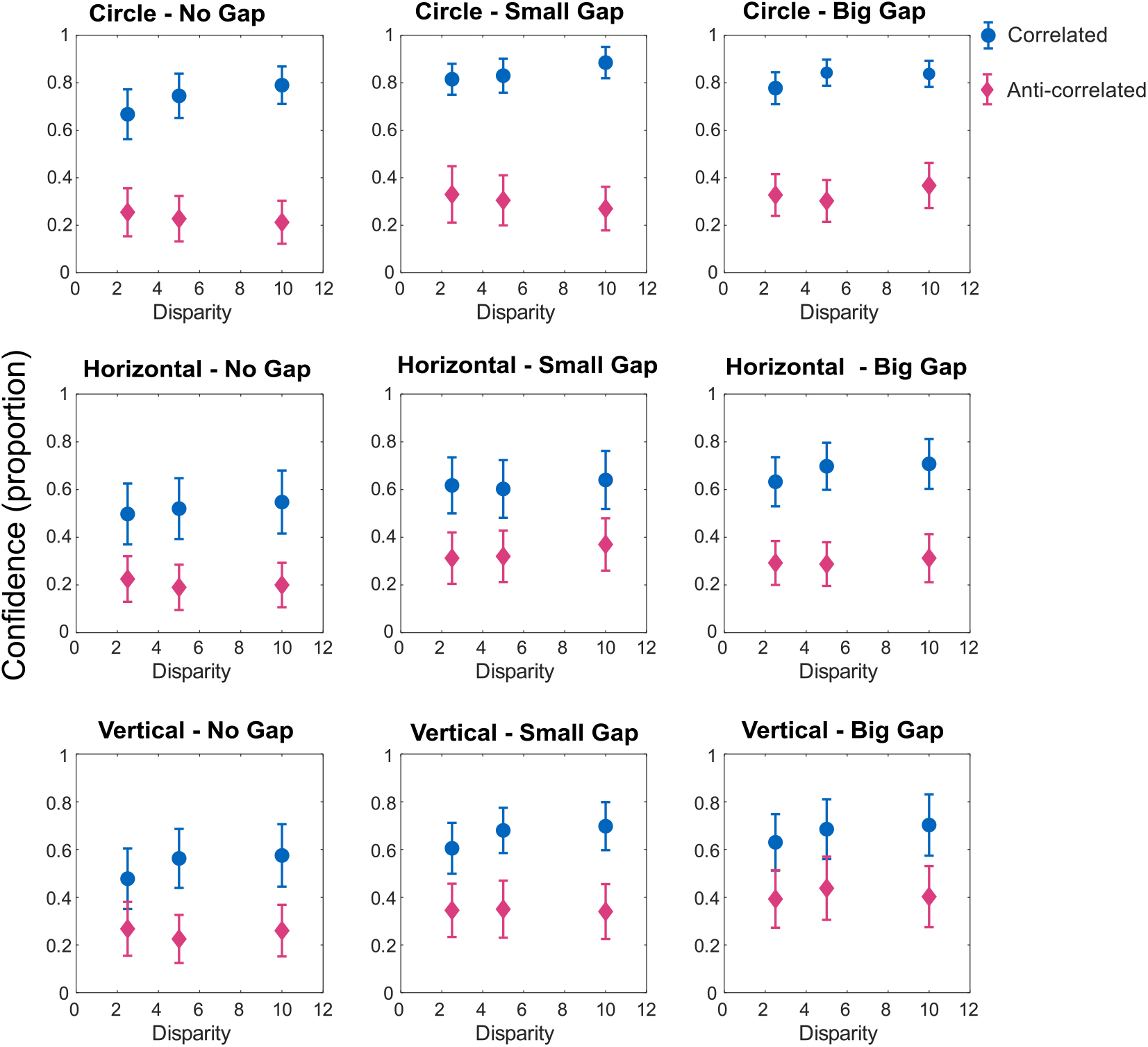
Mean confidence (proportion) for CRDS (blue) and ACRDS (pink) are plotted as a function of disparity and gap size and shape. Error bars show *±*1 standard error of the mean

These results were analysed using a linear mixed effects model, with the shape (circular, horizontal or vertical) and correlation (correlated or anti-correlated) as categorical factors, and separation and disparity as linear covariates. A correlationby-distance interaction term was also included to determine whether separation had a greater effect on confidence for ACRDS than for CRDS. Participant was included as a random factor, with random intercepts and slopes against disparity. The results are summarised in table 2. Confidence ratings were significantly lower for ACRDS than for CRDS. They were also significantly lower for the horizontal and vertical stimuli that for the circular stimuli. Ratings tended to increase with increasing gap size, but were not affected by disparity. There was no significant distance-by-correlation interaction, meaning that the separation did not affect confidence differently for correlation and ACRDS. The effects of correlation, shape and distance are summarised in figure 13.

**Figure 13.**
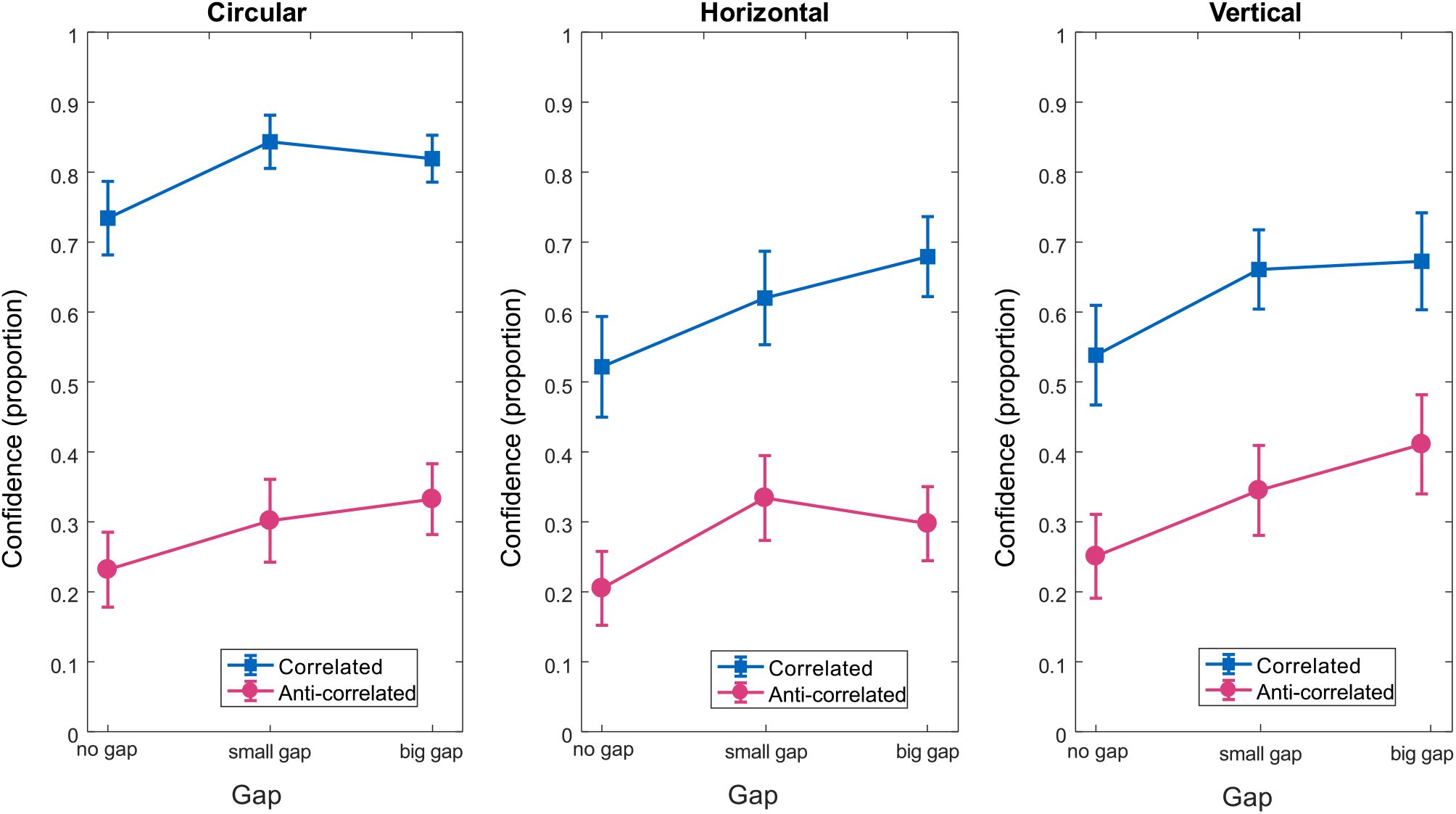
Summary of confidence in depth detection as a function of gap size for CRDS (blue) and ACRDS (pink), and each shape on separate plots. Error bars show *±*1 standard error of the mean

**Table 1.** Estimates of effects from the generalised linear mixed effects regression for the proportion of near versus far responses in CRDS and ACRDS stimuli

| Shape | Correlation | Gap Size | $\beta$ | SE | tStat | DF | pValue |
| --- | --- | --- | --- | --- | --- | --- | --- |
| Circle | CRDS | 0 | 0.05095 | 0.018469 | 2.7586 | 58 | 0.0077524* |
| Circle | CRDS | 19.2 | 0.058554 | 0.020582 | 2.845 | 58 | 0.0061278* |
| Circle | CRDS | 38.4 | 0.061829 | 0.021556 | 2.8682 | 58 | 0.0057471* |
| Circle | ACRDS | 0 | -0.0011257 | 0.0021927 | -0.5134 | 58 | 0.60962 |
| Circle | ACRDS | 19.2 | -7.8727e-05 | 0.0013605 | -0.057867 | 58 | 0.95405 |
| Circle | ACRDS | 38.4 | 0.0027754 | 0.0021185 | 1.3101 | 58 | 0.19532 |
| Horizontal | CRDS | 0 | 0.042448 | 0.015851 | 2.6779 | 58 | 0.0096197* |
| Horizontal | CRDS | 19.2 | 0.038207 | 0.015074 | 2.5347 | 58 | 0.013976* |
| Horizontal | CRDS | 38.4 | 0.039287 | 0.015882 | 2.4736 | 58 | 0.016324* |
| Horizontal | ACRDS | 0 | 0.004145 | 0.0030626 | 1.3534 | 58 | 0.18117 |
| Horizontal | ACRDS | 19.2 | 0.0069563 | 0.0029483 | 2.3594 | 58 | 0.021692* |
| Horizontal | ACRDS | 38.4 | 0.0045466 | 0.0033093 | 1.3739 | 58 | 0.17476 |
| Vertical | CRDS | 0 | 0.023867 | 0.011142 | 2.1421 | 58 | 0.036395* |
| Vertical | CRDS | 19.2 | 0.03677 | 0.014867 | 2.4733 | 58 | 0.016339* |
| Vertical | CRDS | 38.4 | 0.048554 | 0.017404 | 2.7899 | 58 | 0.0071229* |
| Vertical | ACRDS | 0 | 0.0018335 | 0.0018846 | 0.97288 | 58 | 0.33465 |
| Vertical | ACRDS | 19.2 | 0.0050179 | 0.0022086 | 2.2719 | 58 | 0.026817* |
| Vertical | ACRDS | 38.4 | 0.007975 | 0.0036747 | 2.1702 | 58 | 0.034096* |

\*indicates significant p value.

**Table 2.** Estimates of effects from the linear mixed effects regression for confidence judgements in CRDS and ACRDS stimuli

| Shape | $\beta$ | SE | tStat | DF | pValue | CI | CI |
| --- | --- | --- | --- | --- | --- | --- | --- |
| Circle | 23.162 | 3.7038 | 6.2536 | 533 | 8.2323e-10 | 15.886 | 30.438 |
| Horizontal | -4.0333 | 1.0966 | -3.678 | 533 | 0.00025894* | -6.1876 | -1.8791 |
| Vertical | -2.5611 | 1.0966 | -2.3355 | 533 | 0.019889* | -4.7153 | -0.40689 |
| Correlation | -14.711 | 2.369 | -6.2099 | 533 | 1.0675e-09* | -19.365 | -10.057 |
| Gap Size | 2.5111 | 0.93709 | 2.6797 | 533 | 0.0075966* | 0.67028 | 4.3519 |
| Disparity | 0.083911 | 0.065264 | 1.2857 | 533 | 0.1991 | -0.044296 | 0.21212 |
| Correlation x Gap | -0.15556 | 1.0966 | -0.14185 | 533 | 0.88725 | -2.3098 | 1.9987 |

\*indicates significant p value.

#### Relationship between Confidence and Performance

For each stimulus, we recorded a near/far depth judgement and an indication of whether or not the participant felt confident in this judgement. Figure 14 plots heatmaps of the relationship between confidence and performance, for CRDS and ACRDS, summed over all participants, disparities, separations and stimulus types. Performance was summarised at the number of consistent responses such that, for example, 20 near responses out of 20, and 20 far responses, were both coded as 100% consistent, while a stimulus for which there were 10 near and 10 far responses was coded as 50% consistent. This consistency measure does not depend on whether responses are in the forward or reversed direction, relative to the stimulus disparity. For CRDS, results are clustered in the top-right corner, indicating generally high levels of both consistency and confidence. For ACRDS, confidence was generally low, consistent with a low level of consistency in responses.

**Figure 14.**
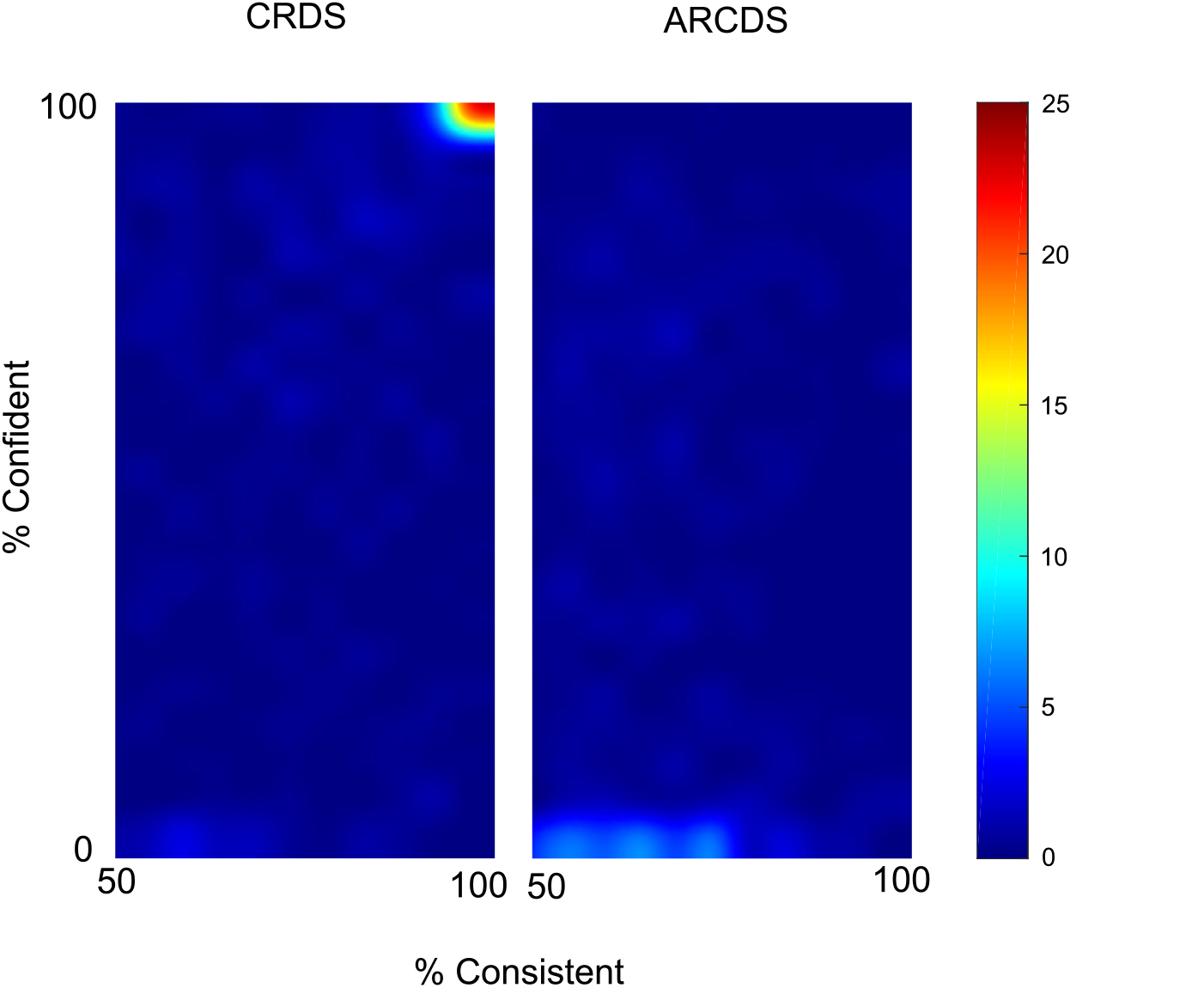
Relationship between consistency of, and confidence in, responses, for CRDS (left) and ACRDS (right). Colour indicates the percentage of responses, for each trial, at each level of consistency and confidence, averaged across all participants and stimuli. High levels of both consistency and confidence were recorded for CRDS, and low levels of both measures for ACRDS.

### Discussion

The aim of our psychophysical experiment was to understand how the spatial separation (the size of the gap) and overlapping correlated and anti-correlated elements influences our perception of depth in ACRDS.

Predictably, our results show forward depth perception for all CRDS stimuli. Depth perception for horizontal and vertical stimuli was not influenced by the size of the gap or type of stimulus (circle, horizontal or vertical). In addition for CRDS, confidence increased with gap size. Confidence was highest for circular stimuli and lower for horizontal and vertical stimuli.

In contrast, for ACRDS confidence in judgements was at or below 50% for all conditions. These are significantly lower than the respective confidence ratings for CRDS. Furthermore, depth perception for ACRDS was variable depending on the type of stimulus.

For circular ACRDS there was, on average, no depth perception for any gap size. However, reversed depth as predicted by Aoki et al.^17^ was seen by some participants in the no-gap condition. For the small-gap, there was no depth discrimination shown by any participants, and for the big-gap only two participants reported forward depth.

For horizontal stimuli, on average there was a tendency towards forward depth for ACRDS with a small-gap, but no forward or reverse perception of depth for stimuli with no-gap or a small-gap. While the majority of participants reported no depth, a minority reported forward depth, and one participant reported reversed depth.

Finally, for vertical stimuli, on average depth was perceived in the forward direction for the small- and large-gap ACRDS, while there was no perception of depth with the no-gap stimuli. This trend reflected the forward-depth perception in a minority of participants.

Aoki et al.^17^ found that removing the gap between the target and the surround for ACRDS increased the perception of reversed depth. However the removal in the gap results in the potential for overlap between the anti-correlated dots in the target and the correlated dots in the surround. This results in decorrelation (an average correlation of zero) at the edges on opposite sides of the target in each eye. This occlusion occurs naturally in the phenomenon of da Vinci stereopsis when the edges of vertically oriented stimuli occlude the surface from the opposite eye^50^ as illustrated in figure 5. If these uncorrelated regions were taken to indicate the presence of half-occlusions, we would predict that depth would be perceived in the reverse direction.

Results indicate that there was little evidence of reversed depth for the vertical edges, and one condition (small-gap) showed a significant tendency for depth to be perceived in the forward direction. There was only one participant who reported reversed depth for the horizontal no-gap condition.

This study found no robust evidence for reversed depth perception in any condition, supporting the findings by Hibbard et al.^19^. The anti-correlated circle condition with no-gap produced the highest proportion of reversed depth perception, consistent with previous results^47^. Furthermore the circle conditions, no-gap and small-gap, were the only conditions not reporting any forward depth perception. The trend towards depth perception in the forward direction, while not consistent with the information provided by first-order disparity channels, is predicted by the depth signalled by second-order mechanisms. The conflicting depth signalled by first- and second-order mechanisms for ACRDS is explored in the following section.

## First- and Second-order responses to correlated and anti-correlated random dot stereograms

The perception of depth depends on multiple mechanisms, including first- and second-order channels, tuned to a variety of scales and orientations. How these contribute to the perception of depth depends on how the information they provide is processed in higher cortical areas. In order to understand the contribution of first- and second-order mechanisms to perceived depth in correlated and ACRDS, we modelled the responses of these mechanisms to our stimuli.

### Methods

#### First- and Second-Order Mechanisms

First-order disparity-sensitive mechanisms were implemented using a standard binocular energy model. The first stage of this model was a quadrature pair of Gabor filters:

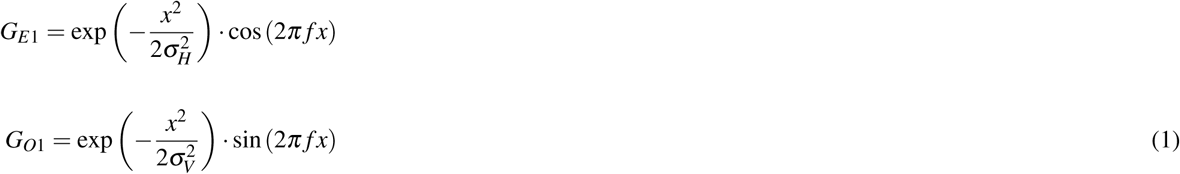

where (*x, y*) is the spatial position, *f* is the spatial frequency and *σ_H_* and *σ_V_* determine the horizontal and vertical extent of the Gaussian envelope. All filters were tuned to a vertical orientation, and spatial frequencies of 4, 8 and 16 cycles /degree were used. The values of *σ_H_* and *σ_V_* depended on spatial frequency as follows:

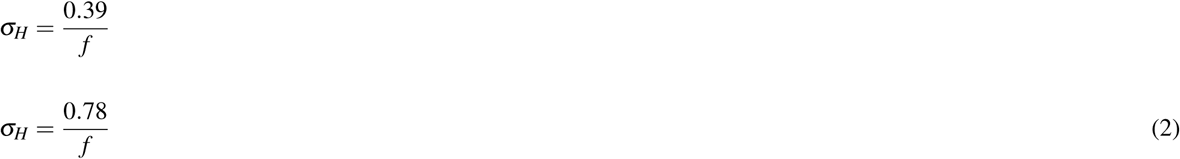

The left and right images were both convolved with each filter, to produce four responses:

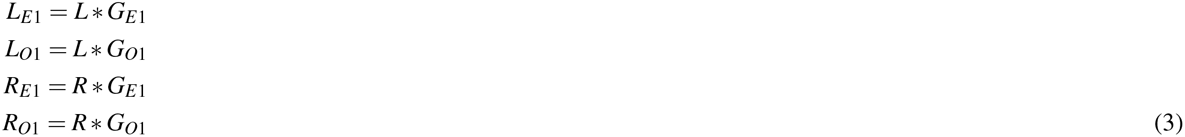

Binocular energy was calculated as:

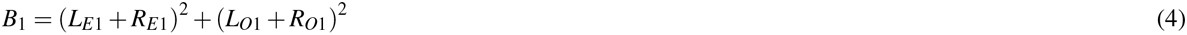

The second-order mechanisms included two filtering stages. The first used the same filters as first-order mechanisms. Left and right monocular energy responses were calculated as:

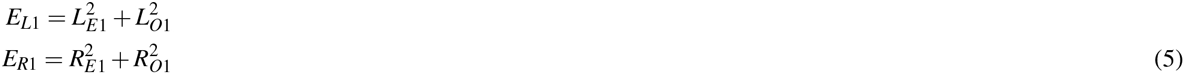

Second-order filters (*G_E_*_2_*, G_O_*_2_) had the same shape as first-order filters, but a frequency tuning of 0.8 cycles/degree. Monocular responses were then calculated by convolving the left and right energy responses with the second-order filters,

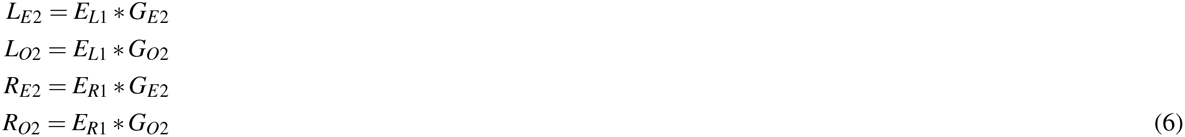

and combining these to produce the second-order energy response:

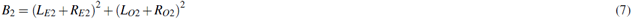

#### Stimuli

Responses to stimuli with the same random dot pattern as that used in the psychophysical experiments were analysed. In each stimulus, the upper half was correlated and presented with zero disparity, and the lower half either correlated or anti-correlated and presented with a disparity of 10 arc min.

#### Procedure

First- and second-order binocular energy responses were calculated for 1000 samples. In all cases, the values used to calculate the energy response were taken from the central row of the left image, and from a row in the right image corresponding to a horizontal disparity of ±30 arc min, sampled at intervals of 1 arc min. All convolutions were calculated by multiplication in the Fourier frequency domain.

### Results

Figure 15 shows the mean, over 1000 trials, of the first order energy response, as a function of vertical position and horizontal disparity. Results are shown separately for each of the three spatial frequencies. Results are plotted separately for CRDS and ACRDS. Results were normalised for each frequency by dividing by the maximum of the mean response. The horizontal lines show the target- and surround regions, with a non-zero and zero disparity, for correlated (blue) and anticorrelated (pink) regions. For CRDS, there is a peak in response for all frequencies at the correct disparity, flanked by a minimum in response, the disparity of which depends on the frequency tuning of the filter. This pattern reflects the tuning function shown in figure 1. For the ACRDS, this response is reversed, such that there is a minimum response at the correct disparity. Reversed depth has been attributed to evidence against the presence of stimulus at the correct disparity,^17^ or by the flanking false peaks.^21^

**Figure 15.**
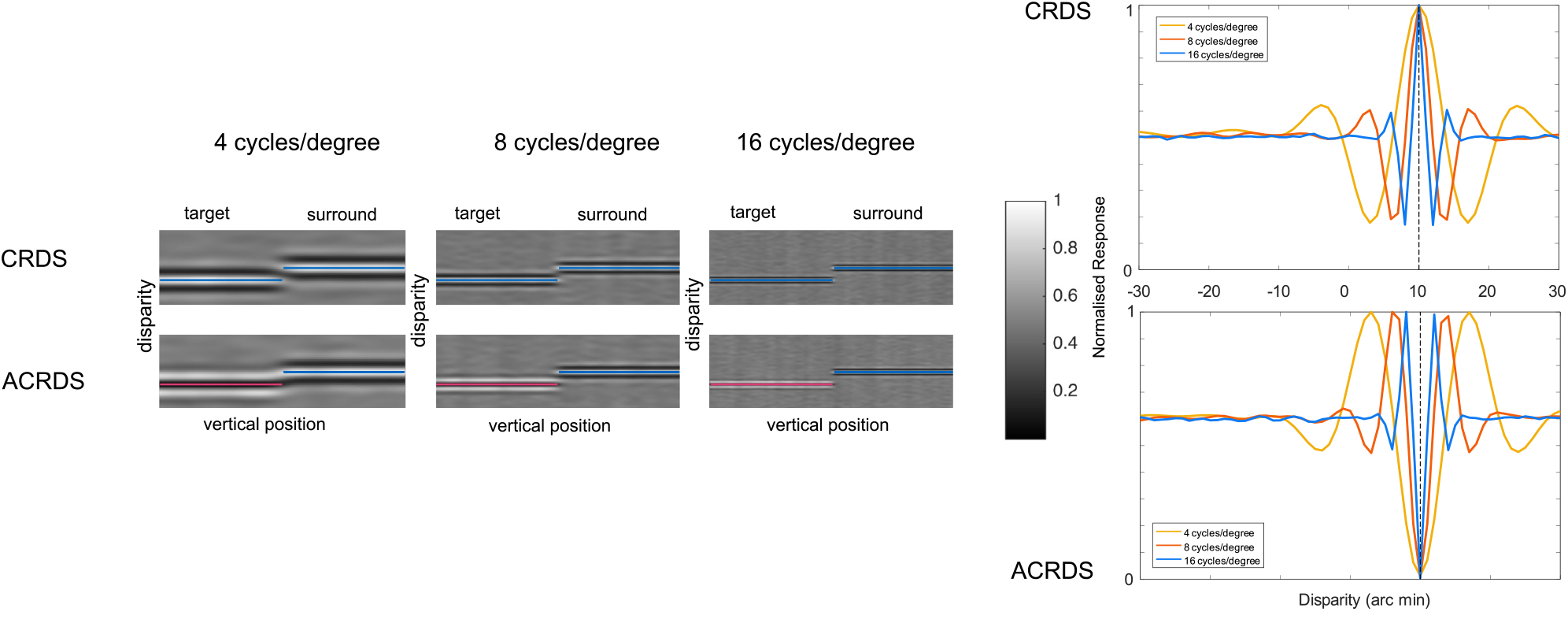
The responses of first-order mechanisms, tuned to 4, 8 and 16 cycles/degree, to CRDS (top row) and ACRDS (middle row). In each image, horizontal and vertical position represent the spatial location and preferred disparity of the filters, respectively, and pixel brightness the strength of response. The target (correlated or anticorrelated) is on the left of each image and the zero-disparity surround is on the right. The disparities of these are indicated by the horizontal lines (blue for correlated regions, pink for anti-correlated regions). The two graphs on the right show the disparity tuning, averaged over the target regions, for all three frequencies, for CRDS (top) and ACRDS (bottom).

Figure 16 shows the mean of the second-order energy response, plotted in the same way as for the first-order response. For CRDS, these results show a very similar pattern to the first-order response, except with much coarser spatial resolution, due to the low-frequency tuning of the second-order filters. For the ACRDS, the inversion of the disparity tuning function is absent. This means that, for ACRDS, second-order filters provide a response that is consistent with the forward disparity, and consistent across the three spatial frequencies of first-order filtering.

**Figure 16.**
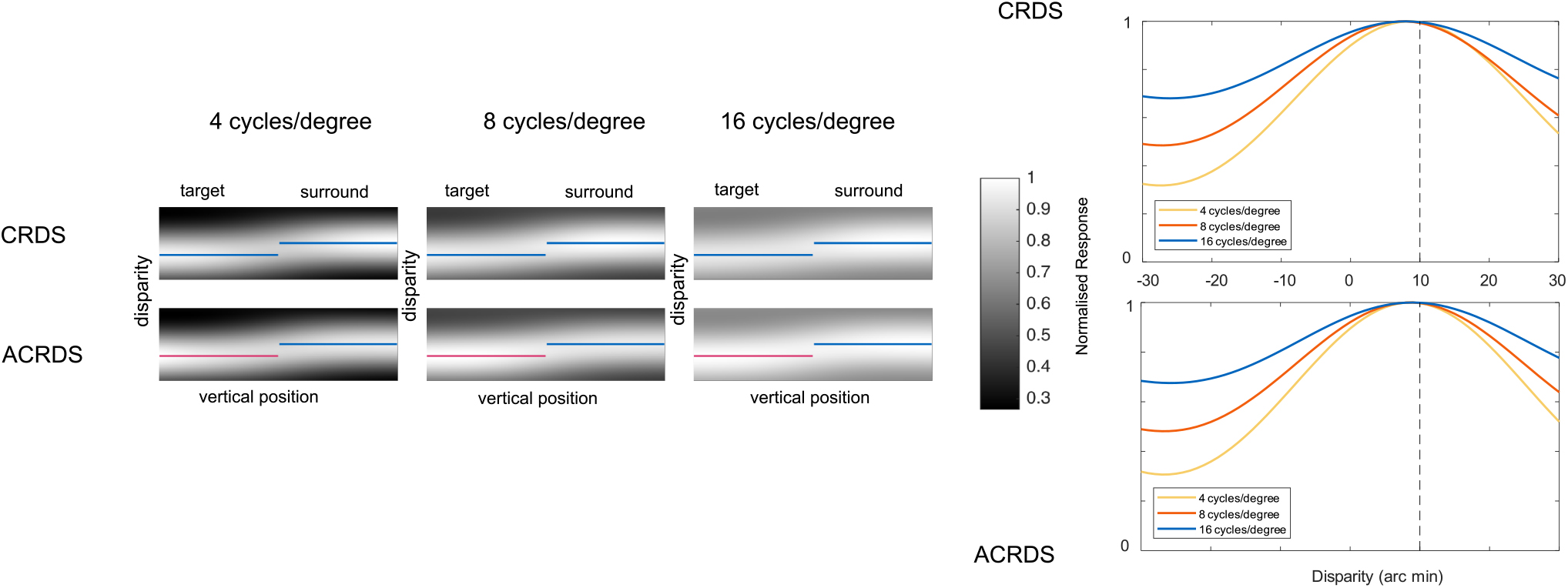
The responses of second-order mechanisms, with first-stage filters tuned to 4, 8 and 16 cycles/degree, to CRDS (top row) and ACRDS (middle row). The second stage filters are in all cases tuned to 0.8 cycles/degree. Results are plotted in the same way as for first-order mechanisms (Figure 15).

### Discussion

The purpose of the second study was to investigate first- and second-order contributions to the perception of depth. This analysis focused on modelling responses to the stimuli used in this study. The reversal of apparent depth found for some ACRDS has been linked to the inversion of the disparity tuning function of binocular cortical cells. This inversion of disparity tuning does not occur for second-order mechanisms, due to the rectifying non-linearity that occurs between the first- and second-stage filtering. This means that second-order units indicate depth in the forward direction for both CRDS and ACRDS.

Hibbard et al. (2016) proposed that second-order mechanisms may provide some error correction to the first-order disparity estimations where regions are decorrelated^15^. This is consistent with the perception of forward-depth in ACRDS shown here, despite the lack of a clear binocular cross-correlation peak at the target disparity. While the first- and second-order responses have been shown to be highly correlated in natural stimuli, second-order mechanisms can increase the accuracy of depth perception, acting as a ‘back-up’ mechanism when first-order mechanisms do not provide a reliable signal.^31^

The existence of both first- and second-order mechanisms substantially complicates the attempt to link depth perception in ACRDS to neural population responses. It may also accounts for why there is so much variability across individual participants.^15^ As well as providing an impoverished, inconsistent signal across scales in the first-order channels, conflicting estimates can arise between first- and second-order channels.

### General Discussion

Our findings suggest that neither particular spatial configurations of stimuli, nor gap sizes, contributed to robust perception of reversed depth in ACRDS. As expected, CRDS stimuli were all perceived with depth in the geometrically predicted forward direction. For ACRDS, most participants reported no depth at all, and in some cases there were significant positive slopes consistent with forward depth perception for the vertical (small-gap & big-gap) and horizontal conditions (small-gap only). For the anti-correlated circular conditions, overall, there was very little perception of forward depth for any gap size, with the exception of 2 participants when the gap was at its biggest. For the no-gap condition reversed depth was reported by 4 participants.

In order to better understand the contributions of the first- and second-order disparity processing stages to the perception of depth, we modelled the responses to the stimuli used in this experiment using a standard binocular energy model, adapted to include second-order filters.^23^ Results from the first-order energy response show a peak for CRDS at the correct disparity, as would be expected. For ACRDS, this peak is reversed. The second-order energy response shows a similar peak location as the first-order response for CRDS, but with much broader disparity tuning. However, for the ACRDS there was no inverse peak, as predicted by the rectifying non-linearity occurring prior to the second stage of filtering. This indicates that second-order mechanisms signal forward depth for both CRDS and ACRDS.

The conflicting signals from the first- and second-order channels may explain the contradictory results between studies^17, 19^ and between our conditions, and participants. Wilcox and Allision (2009)^31^ proposed that the second-order mechanism provides a back up to the first-order mechanism. When there is a reliable luminance-based disparity signal then the first-order mechanism will be relied upon. However, when there is an ambiguous signal, such as when using anti-correlated or uncorrelated dots, second-order signals are used.^31^ Strong evidence to support this is found in individuals with strabismus,^56^ a condition where the eyes are abnormally aligned. Standard clinical tests suggest that individuals with strabismus are stereo-deficient, however McColl et al. (2000) demonstrated that some individuals were still able to perceive depth in their task. Wilcox and Allision (2009) suggest this depth perception was via coarse disparity signals from the second-order mechanisms.

A possible explanation for our results may lie in the depth cue combination model proposed by Landy et al.^57^. Depth cues will vary throughout a scene, and while each individual cue contributes to the estimated weighted average of depth, these will vary in the quality of information they contain.^57^ Where a cue does not provide a depth estimate, provides an unreliable estimate, or one that contradicts the other cues, it will be down-weighted. Landy et al. suggest a mechanism of combination that statistically estimates depth from weighted averages which depend on the interactions^57^ and correlations^15, 58^ between responses. While their model describes cues in terms of qualities of the stimulus, such as texture, shading and parallax, we have shown that, even within a cue, conflicts might exist, such as between estimates from first- and second-order mechanisms. Therefore it is likely that the average weighted estimate includes the error checking response of the second-order mechanism before interpreting depth. While it is still unknown whether first- and second-order mechanisms co-exist or transition from one to another,^31^ our results suggest that when first- and second-order responses are in agreement, depth is perceived normally. Conversely, when the responses from first- and second-order mechanisms conflict, depth would be expected to be perceived at the stage consistent with the direction indicated by the more reliable channel, with the highest decision weight. In the vertical and horizontal stimuli this was the second-order mechanism. However, the spatial complexity of the circular condition suggests that neither first-nor second-order mechanisms are reliable, and therefore the visual system was unable to unambiguously reconcile depth. This would reflect the inability of the second-order channel to reliably signal depth other than in stimuli with very simple spatial variations in surface layout.^42^

Reversed depth perception is not routinely found in centrally presented targets^5, 19^ however, our results are consistent with a recent paper^59^ that found reversed perception of ACRDS in the periphery. Zhaoping and Ackermann (2018)^59^ account for their findings through reduced (or absent) feedback connections in the periphery, resulting in a reduction of verification for stimulus features (see Zhaoping (2017)^59^ for a detailed discussion of the feedforward-feedback-verify-weight network model). An important observation to make is that the stimuli used in our study were neither strictly centrally nor peripherally presented, but ‘mid-way’ between the two positions used by Zhaoping and Ackerman. Since second-order channels are also strongest in central fields, and weaker or absent in the periphery, this may also account for these findings. Therefore, it is also likely that the reversed perception in this condition (circular no-gap) for some participants, was a result of the combination of the part-peripherally located presentation and the complexity of the stimulus.

Since spatially separating the reference and the surround has been found to reduce stereoacuity,^17, 46^ it has been suggested that the spatial gap may be the responsible for the lack of robust reversed depth perception in ACRDS. Consistent with Kamihirata et al.^47^ we found reversed depth when there was no gap, which diminished as the gap size increased. However, neither the horizontal nor vertical conditions showed this trend. This implies that the absence of the gap is not exclusively responsible for promoting the perception of reversed depth. The relatively complicated shape of the circle and its annulus when there is no-gap and thus the opportunity for overlap also plays a role in reversed depth, through reducing the effectiveness of the second-order channel to signal forward depth. Cumming et al.^12^ found that reversed depth perception from ACRDS reduced as a function of increased dot density. Perception of reversed depth has been found at low dot densities (between 1% and 11%).^12, 18^ At high dot densities it is more likely that there is overlap between the dots in each eye, increasing the chance of an ambiguous signal.

A clear finding arising from ACRDS studies^12, 17, 19^ is that there is considerable variability between individuals’ reported perception of depth. Recent findings for robust reverse perception of ACRDS in the periphery^59^ suggest that ACRDS still has an important role to play in allowing us to understand how the visual system learns to process depth. Cumming et al.^12^ attempted to improve depth perception for ACRDS using training with feedback, but were unsuccessful. They used trial by trial feedback with over 10 000 presentations of ACRDS, yet performance did not improve. Recent advances in perceptual learning research indicates that crucial factors in learning is individual confidence and task difficulty.^60–62^ Our results show a very clear difference between participants’ confidence in their judgements for CRDS and ACRDS. Perceptual learning to improve depth perception in anti-correlated stimuli may benefit from these advancements by focusing on improving participant confidence.

### Conclusion

How the visual system combines monocular sensory information into a binocular percept is an important question in vision research. Based on the binocular energy response from first-order mechanisms, it has been proposed that reversed depth perception should occur when presenting an ACRDS target with a CRDS surround. However, these predictions have only taken into account the first-order responses, ignoring the squaring non-linearity of the error checking,^15^ ‘back up’ function^31^ of the second-order mechanism. Our results are consistent with an increasing number of studies that show second-order summation occurs as an independent process taking place after first-order processing, in motion,^63, 64^ perceived contrast^65^ and binocular phase^66^ and stereopsis.^40^. Finally, while depth perception for ACRDS was mostly ambiguous, the positive slopes found in some conditions indicate a trend for perception in the forward direction, rather than reversed depth. Furthermore, confidence for ACRDS was poor in every condition, and increasing participant confidence may be the key to improving reversed depth perception.

## Author contributions statement

J.A and P.H. conceived the experiment; J.A. conducted the experiment; J.A. and P.H. analysed the data; P.H. derived the models; J.A and P.H. wrote and reviewed the manuscript.

## Additional information

The authors declare no competing interests.

